# H3.1^Cys96^ oxidation by mitochondrial ROS promotes chromatin remodeling, breast cancer progression to metastasis and multi-drug resistance

**DOI:** 10.1101/2022.12.01.517361

**Authors:** Flavio R. Palma, Fernando T. Ogata, Diego R. Coelho, Kirthi Pulakanti, Alison Meyer, Yunping Huang, Jeanne M. Danes, Matthew J. Schipma, Cristina M. Furdui, Douglas R. Spitz, Benjamin N. Gantner, Sridhar Rao, Vadim Backman, Marcelo G. Bonini

## Abstract

Transcription stability enforces cellular identity and is tightly controlled by restrictions imposed on both transcription factor function and target gene accessibility. Progression of cancer to metastasis and multi-drug resistance requires fluid transcriptional programs that can explore different genomic landscapes to enable clonal expansion of aggressive and treatment resistant phenotypes. Here, we show that increased levels of H_2_O_2_ produced in mitochondria leads to H3.1 oxidation at Cys96, a distinctive redox sensitive amino acid residue restricted to this histone variant, in the nucleus. The oxidation of Cys96 promotes the eviction of H3.1 from chromatin and its exchange with H3.3, thereby opening silenced portions of the chromatin. Mutation of Cys96 by an oxidation-resistant serine residue or quenching nuclear H_2_O_2_ reversed chemotherapy resistance and drove established metastatic disease into remission. Together, these results show that increased mitochondrial H_2_O_2_ production, characteristic of metabolic dysfunction, promotes transcriptional plasticity by removing structural chromatin restrictions imposed by the redox sensitive histone variant H3.1. We suggest that this new regulatory nexus between cancer metabolism and chromatin remodeling controls chromatin states that enable cancer progression and drug resistance acquisition.

## INTRODUCTION

Most of the trillions of cells in the human body remain restricted to a narrow range of phenotypic states necessary to maintain function and organismal health. The acquisition of inappropriate phenotypes leads to disease, as best exemplified by cancer and metastasis^1^. Lineage-specific gene transcription, a critical mechanism for preserving cellular identity, is enforced through limits on the activity of transcription factors^2-4^ and limits imposed by the chromatin architecture itself^5,6^. These latter epigenetic mechanisms that broadly influence chromatin organization occur largely through post-translational modifications of highly conserved histone proteins^7,8^ that together form the minimal structural unit of chromatin, the nucleosome. In addition to these modifications, exchange of core histone proteins with different histone variants also plays an important transcription regulatory role^9-11^.

As an example, the human histone variant H3.3 is enriched over H3.1 in nucleosomes undergoing active transcription^10^, while H3.1 marks silent chromatin^12^. Accumulating evidence supports the concept that the balance between H3.1 and H3.3 occupancy has important implications for cellular gene expression, identity, and function^9,13,14^. Despite this, the mechanisms that control histone exchange are not completely understood. The modest differences in primary structure observed in histone proteins have permitted phylogenetic analysis supporting a role for H3.3 as the source of all H3 variants in animals and fungi, while H3.2-like alleles appear to be in all animal genomes^15^. In contrast, canonical H3.1, notable for the acquisition of a cysteine residue at position 96, is restricted largely to mammals^15^. While amino acid residue 96 is positioned within the histone “core”, structural modeling studies suggest that it is likely accessible and could even accommodate post-translational modifications without destabilizing the histone structure^16^. Given the unique spatial location of this amino acid position within the nucleosome and the redox-sensitive nature of the cysteine thiol, we examined whether the local oxidative context could regulate the oxidative state of H3.1^Cys96^ and impact chromatin structure.

These studies were stimulated by three key findings. First, the detection of increased concentrations of ROS in the nucleus of breast cancer cells with more aggressive phenotypes. Second, v-Src transformation of MCF10A mammary epithelial cells increased steady state levels of nuclear ROS (nROS) upon activation. Finally, use of a synthetic redox signaling system^17^ engineered to restrict the production of H_2_O_2_ to the nucleus revealed a role for nuclear H_2_O_2_ (nH_2_O_2_) in promoting loss of cellular identity (epithelial to mesenchymal transition - EMT), tumor growth, and metastasis. All of this was dependent on H_2_O_2_ present at levels that failed to trigger cellular stress and DNA damage responses (DDR). Using this system as well as independent orthogonal models, we present data indicating that the local oxidative landscape of the nucleus can shape epigenetic control of cellular identity via promoting the eviction of histone H3.1 and its replacement with H3.3. Critically, we show that this exchange functionality requires Cys96 on H3.1, since mutation of this residue to serine blocks the oxidation-dependent removal of H3.1. Taken together, these data support the concept that H3.1^Cys96^ evolved in mammals to couple nuclear redox state to epigenetic control, and that cancer cells may have learned to exploit this mechanism to survive in the harsh tumor microenvironment and even possibly acquire more aggressive phenotypes. The identification of this novel regulatory mechanism linking metabolic reprogramming to nuclear chromatin structural remodeling suggests that there may be opportunities to target ROS in specific contexts and organelles for therapeutic gain.

## RESULTS

### Steady state nuclear ROS increase with tumorigenicity and metastatic potentials of breast cancer cells

We began testing our hypothesis that alterations in the levels of nROS promote more aggressive cancer cell phenotypes by measuring steady state levels of nH_2_O_2_ in breast cancer cell lines with varying tumorigenicity and metastatic capacity. To accurately measure H_2_O_2_, we used a modified version of the ratiometric redox active biosensor Orp1-roGFP2^18^ designed for nuclear accumulation via fusion with the c-myc nuclear localization sequence (NLS). NLS-Orp1-roGPF2 shows two clear excitation maxima corresponding to the reduced (λ_ex_ = 488 nM) and oxidized (λ_ex_ = 405 nM) forms (Supplementary Figure 1A). Using this construct, we determined that at baseline non-transformed mammary epithelial cells (MCF10A^ER/vSrc^) had the lowest levels of nH_2_O_2_ compared to tumorigenic cells lines (MCF7, BT474, BT20, MB231 and MCF10A^ER/vSrc^ cells transformed by the activation of the v-Src oncogene via stimulation of its fused estrogen-receptor (ER) ligand domain^19,20^, Supplementary Figure 1B). Among the tested breast cancer cell lines, most tumorigenic and metastatic basal cells (BT20, MB231 and transformed MCF10A^ER/vSrc^) had the highest nH_2_O_2_ levels and increased intercellular variability in terms of relative nH_2_O_2_ concentrations (Figures 1A-B) suggesting a link between higher levels of nROS and the activation of transcription associated with the transition of breast cancer towards more aggressive phenotypes that are, typically, less differentiated. Functional studies to determine the effects of nROS required the generation of a system that allowed to controllably and reproducibly generate discrete amounts of H_2_O_2_ confined to the nuclear envelop. For that, we generated clones expressing both NLS-Orp1-roGFP2 and nucleus-targeted D-amino acid oxidase (NLS-DAO), a fungal enzyme that oxidizes D-amino acids simultaneously producing H_2_O_2_ *in situ* (Supplementary Figure 1A)^21^. Because non-transformed MCF10A^ER/vSrc^ cells had the lowest levels of nH_2_O_2_ at baseline, this background was selected to examine the effects of increasing nH_2_O_2_ levels independently of transformation status. Quantification of nH_2_O_2_ with NLS-Orp1-roGFP2 indicated that NLS-DAO expressing MCF10A^ER/vSrc^ cells responded to 10 nM D-Alanine (D-Ala) (Supplementary Figures 1C-D) as expected by increasing nH_2_O_2_. Comparisons with bolus H_2_O_2_ indicated that NLS-Orp1-roGFP2 expressed in MCF10A cells had both dynamic range and a rapid and reversible response to relatively small variations in nROS levels. Experiments were also performed to determine optimal rates of nH_2_O_2_ production that would ensure localization to the nucleus and preservation of DNA integrity to study, to the extent possible, effects that are independent of direct oxidative DNA damage. Figures 1C-E and Supplementary Figure 2 summarizes these studies by indicating that, compared to positive control (500μM H_2_O_2_), 10 nM D-Ala was insufficient to trigger γH2AX accumulation as well as ATM phosphorylation, two sensitive markers of DDR activation (Figure 1C). Additionally, despite elevating nH_2_O_2_ levels (Figure 1D, top panel), 10 nM D-Ala did not significantly increase levels of 8-oxodG (Figure 1D, bottom panel, and Figure 1E) nor elevated levels of DNA double-strand breaks (Supplementary Figures 2A-C), two markers of direct oxidative DNA damage. Finally, our results indicated that there was no measurable increase in amplex red oxidation, a highly sensitive extracellular probe for H_2_O_2_, when D-Ala was used at levels below 10 μM (Supplementary Figure 2D). Therefore, the use of DAO combined with 10 nM added D-Ala to the media was used to identify changes in transcriptional programs activated by nH_2_O_2_ using RNA-seq technology. In Figure 1F, results from comparative transcriptomic analysis between control and cells exposed to the NLS-DAO/10 nM D-Ala system are shown including data collected at an early (4 h) and a later time point (24 h) after activation of nH_2_O_2_ production. Results obtained in the 4 h treatment indicated the upregulation of several genes associated with lineage plasticity acquisition including ALDH1A3, SOX9 (also assessed by RT-QPCR, Figure 1G), members of the Wnt family of transcription factors as well as stem cell transcription factors (POU2F3, KLF4 - verified by RT-qPCR, Figure 1G) and EMT regulating cytokines (TGFα, TGFβ and amphiregulin (AREG), verified by RT-qPCR, Figure 1G). In addition, Gene Set Enrichment Analysis (GSEA) shown in Figure 1H reinforced the idea that a major effect of H_2_O_2_ is to trigger EMT programs given the ranking of representative EMT genes on the diagram. Interestingly, 24 h after exposure to nH_2_O_2_, most of the transcriptomic changes detected in the 4 h treatment were reversed to levels comparable to those observed in control cells at baseline. Taken together, results in Figure 1 indicate that aggressive mesenchymal phenotypes originating from epithelioid breast cancer cells are likely to be sustained, at least in part, by the continuous flow of H_2_O_2_ into the nucleus.

**Figure 1.**
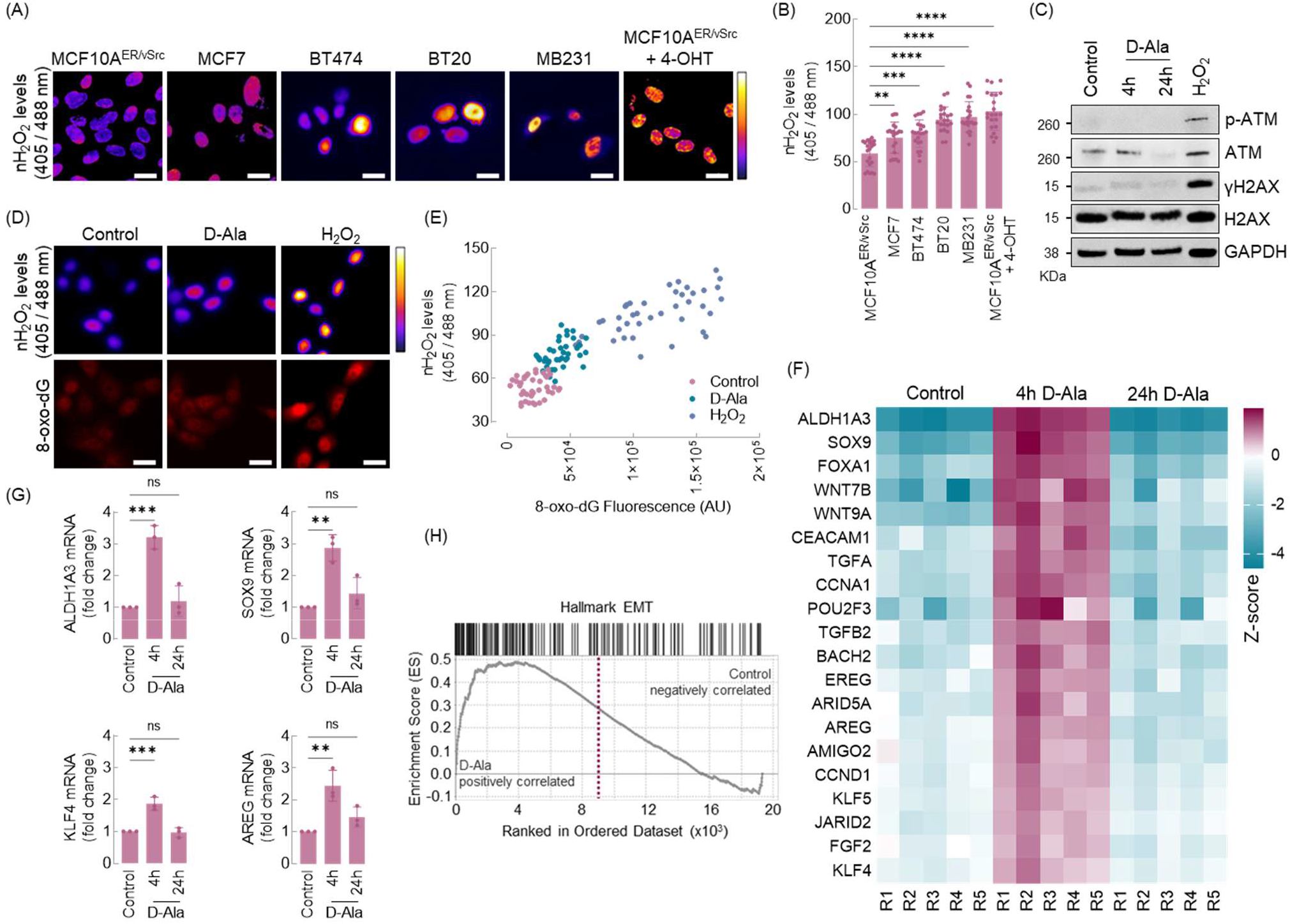
Steady state nuclear ROS increase with the tumorigenicity and metastatic potential of breast cancer cells. (A) The redox state of the nucleus of normal (MCF10A^ER/vSrc^) and cancer cell lines (MCF7, Bt474, BT20, MB231 and transformed MCF10A^ER/vSrc^) was determined using confocal microscopy. Oxidized (λ_ex_ = 405 nM) and reduced (λ_ex_ = 488 nM) roGFP2 signals were acquired and the ratio oxidized/reduced was calculated using ImageJ (shown as a heatmap). White bars represent 10 μm. (B) Quantification of oxidized to reduced ratio of NLS-Orp1-roGFP2 in (A). Statistical significance was determined by One-way ANOVA in combination with Tukey’s test, bars represent mean ± SD. ^**^p < 0.01, ^***^p < 0.001, ^****^p < 0.0001, ns - not significant. (C) Protein levels of p-ATM and γH2AX were assessed by Western blot performed after treatment with 10 nM D-Alanine for 4 h and 24 h and exogenous H_2_O_2_. (D) Immunofluorescence analysis of 8-oxo-dG. Cells treated with 10 nM D-Alanine for 4 h and exogenous H_2_O_2_ for 30 min were stained for 8-oxo-dG (lower panel) and the ratio oxidized/reduced of Orp1-roGFP2 was obtained (upper panel). White bars represent 10 μm. (E) Correlation between fluorescence intensity of 8-oxo-dG and oxidized to reduced ratio of Orp1-roGFP2 in (D). (F) Heatmap of core enriched upregulated genes in the 10nM D-Alanine treatment for 4 h and 24 h (RNA-seq). Z-scores were calculated based on expression of each gene (Log_2_ = 1.5 cutoff). (G) qPCR analysis of ALDH1A3, SOX9, KLF4 and AREG in MCF10A^ER/vSrc^ cells treated with 10 nM D-Alanine for 4 h and 24 h. Statistical significance was determined by One-way ANOVA in combination with Tukey’s test, bars represent mean ± SD. ^**^p < 0.01, ^***^p < 0.001, not significant. (H) GSEA enrichment plot of Hallmark genes associated with EMT and upregulated in the 10 nM D-Alanine 4 h treatment.

**Figure 2.**
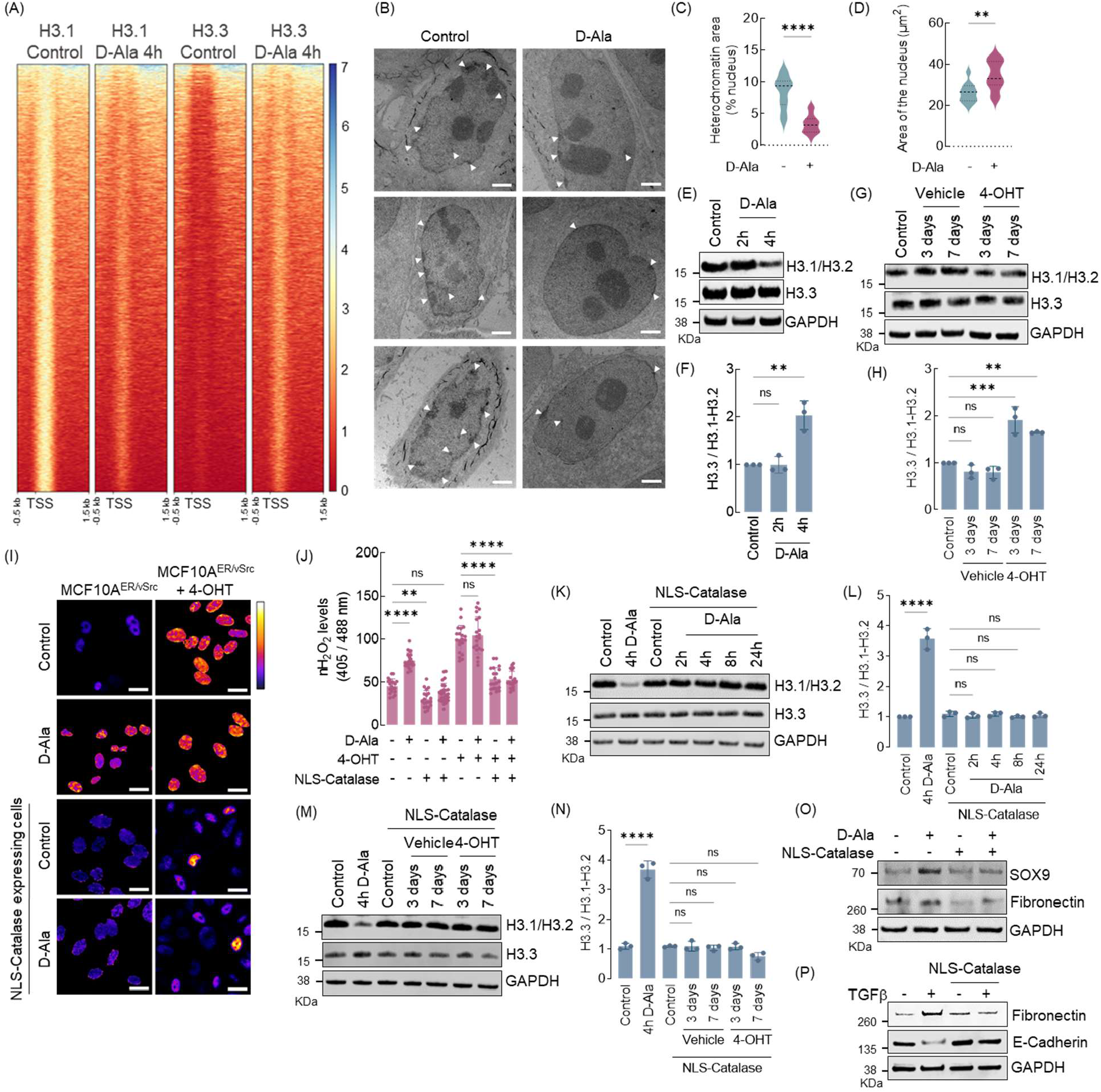
Chromatin remodeling by nuclear ROS involves changes to nucleosome composition. (A) Heatmap of H3.1/H3.2 and H3.3 ChIP-seq peaks across TSS (−0.5kb, +1.5kb) in MCF10A^ER/vSrc^ cells treated with 10 nM D-Alanine for 4 h. (B) Representative images of heterochromatin distribution in cells treated with 10 nM D-Alanine for 4 h assessed by Transmission Electron Microscopy (TEM). White arrows indicate heterochromatin. White bars represent 2 μm. (C) Quantification of perinuclear heterochromatin in (B) using Trainable Weka Segmentation plug-in in Fiji ImageJ. Statistical significance was determined by *t* test. ^****^p < 0.0001. (D) Area of the nucleus of cells in (B). Statistical significance was determined by *t* test. ^**^p < 0.01. (E) Western blot analysis of H3.1-H3.2 and H3.3 in MCF10A^ER/vSrc^ cells treated with 10 nM D-Alanine for 2 h and 4 h. (F) H3.3 / H3.1-H3.2 ratio quantification of (E). Statistical significance was determined by One-way ANOVA in combination with Tukey’s test, bars represent mean ± SD. ^**^p < 0.01, ns - not significant. (G) Western blot analysis of H3.1-H3.2 and H3.3 in MCF10A^ER/vSrc^ transformed cells (3 days and 7 days after treatment with 4-OHT). (H) H3.3 / H3.1-H3.2 ratio quantification of (G). Statistical significance was determined by One-way ANOVA in combination with Tukey’s test, bars represent mean ± SD. ^**^p < 0.01, ^***^p < 0.001, ns - not significant. (I) The redox state of the nucleus of MCF10A^ER/vSrc^ cells expressing NLS-catalase and treated with 10nM D-Alanine or transformed with 4-OHT was determined using confocal microscopy. Oxidized (λ_ex_ = 405 nM) and reduced (λ_ex_ = 488 nM) roGFP2 signals were acquired and the ratio oxidized/reduced was calculated using ImageJ (shown as a heatmap). White bars represent 10 μm. (J) Quantification of oxidized to reduced ratio of NLS-Orp1-roGFP2 in (I). Statistical significance was determined by One-way ANOVA in combination with Tukey’s test, bars represent mean ± SD. ^**^p < 0.01, ^****^p < 0.0001, ns - not significant. (K) Western blot analysis of H3.1-H3.2 and H3.3 in MCF10A^ER/vSrc^ cells expressing NLS-catalase and treated with 10nM D-Alanine for 2 h, 4 h, 8 h and 24 h. (L) H3.3 / H3.1-H3.2 ratio quantification in (K). Statistical significance was determined by One-way ANOVA in combination with Tukey’s test, bars represent mean ± SD. ^****^p < 0.0001, ns - not significant. (M) Western blot analysis of H3.1-H3.2 and H3.3 in MCF10A^ER/vSrc^ transformed cells (3 days and 7 days after treatment with 4-OHT) expressing NLS-catalase. (N) H3.3 / H3.1-H3.2 ratio quantification in (N). Statistical significance was determined by One-way ANOVA in combination with Tukey’s test, bars represent mean ± SD. ^****^p < 0.0001, ns - not significant. (O) Western blot analysis of SOX9 and Fibronectin (EMT markers) in MCF10A^ER/vSrc^ cells expressing NLS-catalase and treated with 10 nM D-Alanine for 8 h. (P) EMT induction of MCF10A cells with TGFβ. Parental and NLS-catalase expressing cells were treated with 10 ng/mL TGFβ for 14 days. The levels of the mesenchymal marker Fibronectin and the epithelial marker E-cadherin were determined by western blot.

### Chromatin remodeling by nuclear ROS involves changes to nucleosome composition

Elegant studies published recently indicated that EMT is preceded by profound chromatin remodeling driven largely by alterations in nucleosome composition. In these studies, Gomes *et al*.^9^ found that the replacement of the H3 histone variant H3.1 by H3.3 was required for TGFβ And TNFα-induced EMT and preceded cancer cell metastasis. H3.1 is unique among H3 histones due to the presence of an oxidizable cysteine residue at position 96. This characteristic, exclusive to H3.1, together with the overrepresentation of EMT-related genes among those upregulated in response to nH_2_O_2_, led us to test the hypothesis that H3.1 oxidation might promote its replacement by H3.3 to trigger EMT. The H_2_O_2_-dependent loss of H3.1 was queried in a series of complementary experiments. First, we performed ChIP-seq experiments using MCF10A^NLS-DAO^ cells. Antibodies specific for H3.1/H3.2 and H3.3 histones were used to immunoprecipitate canonical (H3.1/H3.2) as well as H3.3 variants. Results shown in Figure 2A indicate that nH_2_O_2_ led to significant loss of H3.1/H3.2 canonical histones in nucleosomes close to transcription start sites at 4h after activation of H_2_O_2_ production. The loss of H3.1 was simultaneous to gain of H3.3 at the same time point indicating chromatin decompaction later confirmed using transmission electron microscopy (TEM) (Figures 2B-D). In fact, control MCF10A cells showed a typical pattern of condensed chromatin (H3.1-decorated heterochromatin) deposition around the inner membrane of the nuclear envelope (Figure 2B left panel and Figure 2C). TEM results also show significant dissolution of heterochromatin 4 h after nH_2_O_2_ treatment (Figure 2B right panel and Figure 2C) and an enlargement of the nuclei consistent with chromatin decompaction (Figure 2B lower panel and Figure D). Western blot analysis confirmed loss of H3.1 leading to a net increase in H3.1-H3.2/H3.3 ratio (Figures 2E-F). To investigate if malignant transformation could, similarly to nH_2_O_2,_ induce H3.1/H3.3 exchange, a model of v-Src induced epithelial cell transformation, previously characterized before^19,20^, was used. According to Figures 2G-H, activation of v-Src led to the time-dependent loss H3.1-H3.2 without significant loss of H3.3. This timeline is both consistent with previous studies that showed complete transformation by day 7^19,20^ as well as data shown in Figure 1A indicating that fully transformed MCF10A^ER/vSrc^ cells have higher levels of nH_2_O_2_. In fact, the indispensable role of nH_2_O_2_ both as a mediator of H3.1-H3.2 exchange as well as transformation was confirmed in additional experiments where a nucleus-targeted form of catalase (NLS-catalase) was used. Catalase is an extremely efficient and specific scavenger of H_2_O_2_. According to results shown in Figures 2I-J expression of NLS-catalase in MCF10A^ER/vSrc^ cells prevented the buildup of nH_2_O_2_ in the nucleus during transformation as well as in the 10 nM D-Ala treatment (4 h). Consistent with the idea that H_2_O_2_ is both required for canonical H3 variant destabilization and consequently transformation, expression of NLS-catalase also preserved H3.1-H3.2 both in the context of NLS-DAO/D-Ala induced nuclear H3.1-H3.2 loss (Figures 2K-L) as well as in the case of v-Src-induced transformation (Figures 2M-N). EMT inhibition was confirmed by Western blot analysis of SOX9 and Fibronectin (mesenchymal markers) that were increased 8 h after nH_2_O_2_ in wild type but not NLS-catalase expressing MCF10A cells (Figure 2O). Finally, previous studies showed that EMT induced by TGFβ requires H3.1-H3.2 replacement by H3.3^9^. To determine if H3 variant exchange in response to EMT induction is dependent on an increase in nROS, MCF10A cells and counterparts expressing NLS-catalase were treated with TGFβ (10 ng/mL) for 14 days. Results shown in Figure 2P, indicate that while wild type cells underwent TGFβ-induced EMT (as indicated by the gain of fibronectin and loss of E-cadherin), the transition to a more mesenchymal phenotype was blocked by NLS-catalase expression suggesting that nH_2_O_2_ is needed to license the expression of TGFβ-induced EMT genes via a process that has been shown to involve H3 variant exchange. Together, results shown in Figure 2 indicate that an increase in nH_2_O_2_ levels precedes and directs phenotypic transitions involvement in malignant transformation by facilitating the reconfiguration of nucleosomal H3 variant composition.

### Cys96 oxidation promotes H3 variant exchange and activates EMT gene expression

H3.1 and H3.3 are 97% homologous with Cys96, a redox sensitive residue, being among the only 5 amino acids residues that distinguish them. To examine the hypothesis that the sensitivity of H3.1 to H_2_O_2_-dependent regulation derives from Cys96 oxidation cell lines expressing H3.1 (C96S)-Flag was produced on the MCF10A background. MCF10A^ER/vSrc^ cells expressing both NLS-DAO and H3.1 (C96S)-Flag mutant variant was treated with 10 nM D-Alanine. We found that, unlike the wild type protein, the oxidation-resistant mutant H3.1 (C96S) was no longer responsive to nH_2_O_2_ (Figures 3A-B). These results indicate that a single cysteine residue at position 96 confers sensitivity to ROS-dependent regulation and that the sensitivity to ROS regulation does not dependent on the other 4 unique non-homologous amino acids. To better understand the role of Cys96 in the association of H3 variants with promoter regions of EMT-associated genes, ChIP-qPCR of DNA was performed. We focused the analysis on DNA segments associated with endogenous H3.1 and H3.3 variants as well as the C96S H3.1-mutant. We found that endogenous H3.1 had a decreased association with promoter regions of SOX9, ZEB1, and Fibronectin after treatment with D-Alanine for 8 h, while H3.1^C96S^ mutant showed a slightly increased association with the same promoters at this time point (Figure 3C). Compared to its baseline, endogenous H3.3 showed a H_2_O_2_-dependent increased association with SOX9, ZEB1, and Fibronectin promoters, replacing the endogenous H3.1 after D-Ala treatment. Nuclease-based accessibility assay showed that SOX9, ZEB1, and Fibronectin were more accessible after treatment with D-Ala (4 h) in cells expressing only the endogenous H3.1, while in cells expressing H3.1 (C96S) mutant the accessibility did not change in response to nH_2_O_2_ (Figure 3D). Finally, protein levels of SOX9, ZEB1, and Fibronectin were also increased after treatment with D-Ala in cells expressing only endogenous H3.1, while in cells expressing H3.1^C96S^ mutant levels remained unchanged (Figures 3E-F). Together, these findings show that Cys96 regulates the redox dependent exchange of H3.1 and H3.3 variants via a mechanism that depends on its oxidation and that is required for the expression of genes involved in epithelial to mesenchymal transition.

**Figure 3.**
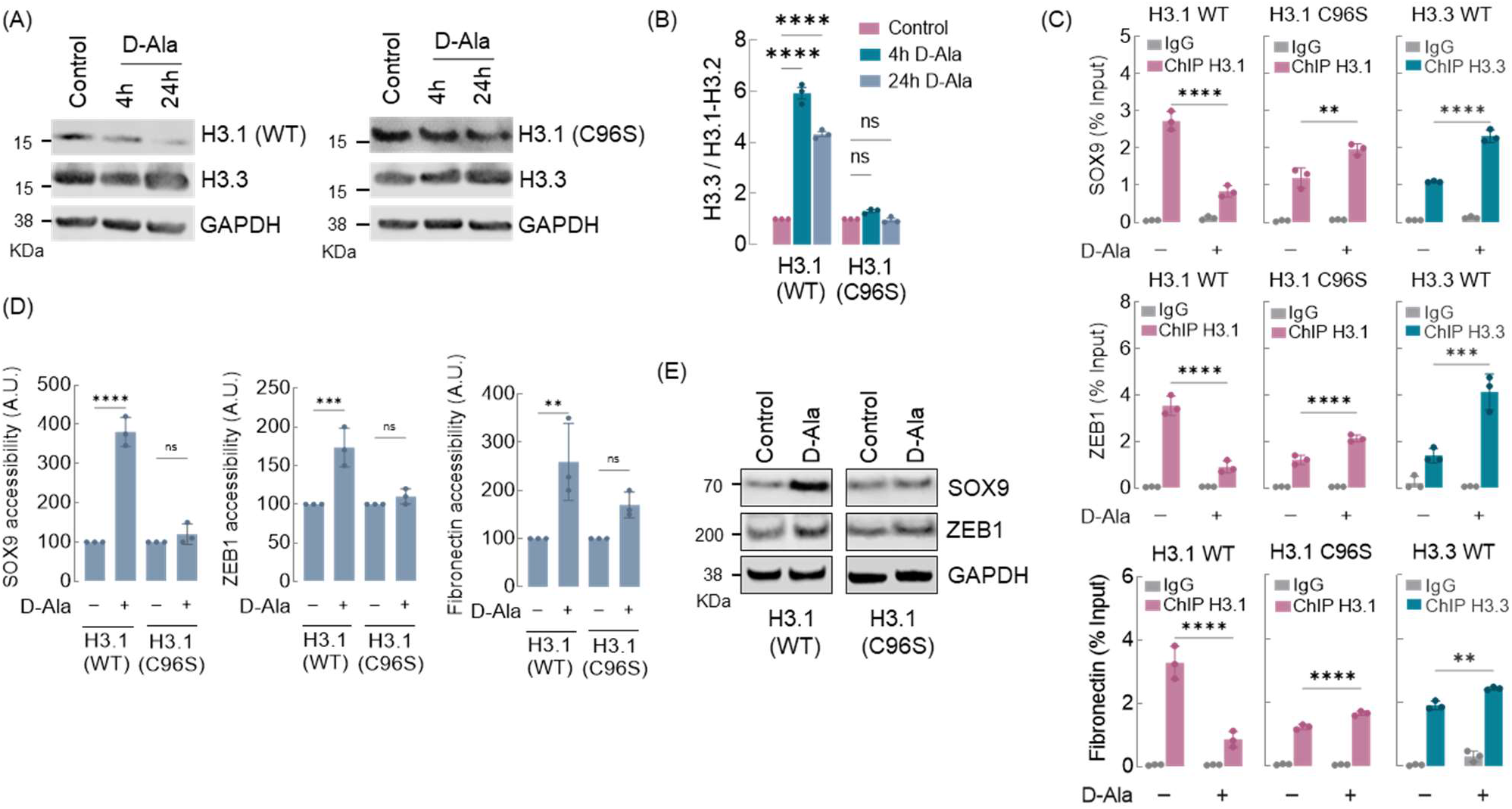
Cys96 oxidation promotes H3 variant exchange and activates EMT gene expression. (A) Western blot analysis of H3.1 (endogenous and C96S) and H3.3 (endogenous) in MCF10A^ER/vSrc^ cells treated with 10 nM D-Alanine for 4 h and 24 h. (B) H3.3 / H3.1-H3.2 ratio quantification of (B). Statistical significance was determined by One-way ANOVA in combination with Tukey’s test, bars represent mean ± SD. ^****^p < 0.0001, ns - not significant. (C) H3.1 (endogenous and C96S) and H3.3 (endogenous) enrichment (ChIP-qPCR) in promoter regions of SOX9, ZEB1, and Fibronectin in MCF10A^ER/vSrc^ cells treated with 10 nM D-Alanine for 8 h. Statistical significance was determined by One-way ANOVA in combination with Tukey’s test, bars represent mean ± SD. ^**^p < 0.01, ^***^p < 0.001, ^****^p < 0.0001. (D) SOX9, ZEB1 and Fibronectin genes accessibility in MCF10A^ER/vSrc^ cells treated with 10 nM D-Alanine for 4 h. Chromatin accessibility was determined in the promoter region of SOX9, ZEB1, and Fibronectin. Statistical significance was determined by One-way ANOVA in combination with Tukey’s test, bars represent mean ± SD. ^**^p < 0.01, ^***^p < 0.001, ^****^p < 0.0001, ns - not significant. (E) Western blot analysis of SOX9 and ZEB1 in MCF10A^ER/vSrc^ cells expressing H3.1 (endogenous and C96S) and treated with 10 nM D-Alanine for 4 h and 24 h.

### nH_2_O_2_ originate from and are regulated by mitochondria

Glycolytic tumors tend to be more aggressive and mesenchymal^22^. In these tumors most of the tumor bioenergenetic needs are supported by metabolic pathways that are largely independent of mitochondria. We hypothesized that another feature of the glycolytic switch that may be advantageous to the evolving tumor is the relieve mitochondria from functioning primarily as a source of ATP to refocus on ROS production needed for oxidative chromatin remodeling. To test this idea, we used MCF10A^ER/vSrc^ cells stably transfected with either nucleus - or mitochondria-targeted Orp1-roGFP2 (mito-Orp1-roGFP2). Transformation induced by v-Src activation significantly increased the levels of nH_2_O_2_ both in mitochondria and in the nucleus (Figures 4A-D). Expression of mitochondria-targeted catalase strongly suppressed H_2_O_2_-driven Orp1-roGFP2 oxidation both in the nucleus and in mitochondria indicating that H_2_O_2_ acting in the nucleus originated from mitochondria. Consistently, mito-catalase expression also suppressed the loss of H3.1 histone in v-Src-transformed cells (Figures 4E-F). In addition, mito-catalase phenocopied the effect of NLS-catalase in suppressing TGFβ-induced EMT as indicated by the inhibition of TGFβ-induced upregulation of fibronectin as well as downregulation of E-cadherin (Figure 4G). Finally, mito-catalase strongly suppressed the expression of nH_2_O_2_-responsive genes in v-Src transformed cells including SOX9, ZEB1 and fibronectin (Figure 4H). Collectively, results in Figure 4 indicate that mitochondria are major sources of H_2_O_2_ involved in chromatin remodeling associated with tumor cell lineage plasticity acquisition and EMT.

**Figure 4.**
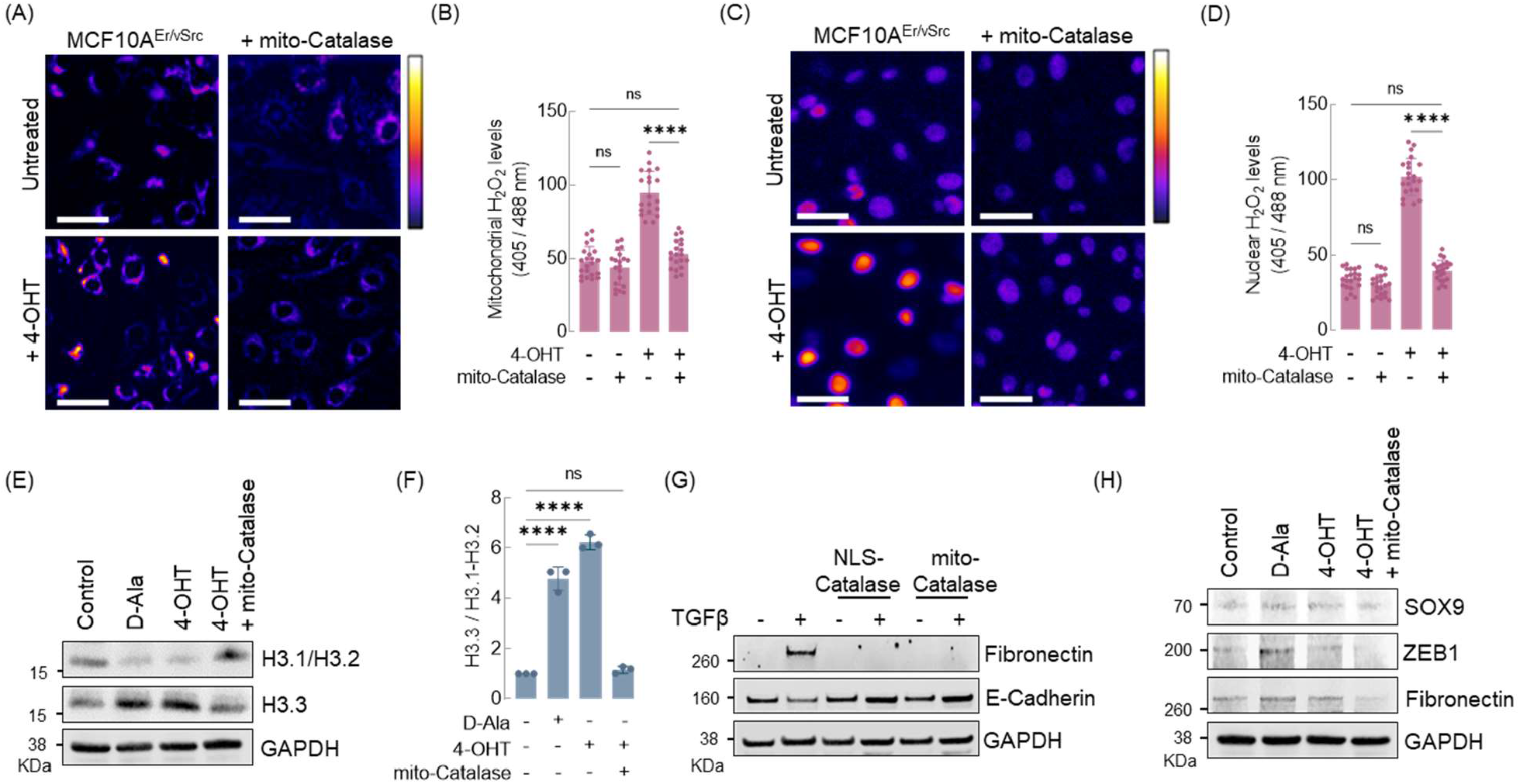
nH_2_O_2_ originate from and are regulated by mitochondria. (A) The redox state of the mitochondria of MCF10A^ER/vSrc^ cells expressing mito-catalase and transformed with 4-OHT (7 days) was determined using fluorescence microscopy. Oxidized (λ_ex_ = 405 nM) and reduced (λ_ex_ = 488 nM) roGFP2 signals were acquired and the ratio oxidized/reduced was calculated using ImageJ (shown as a heatmap). White bars represent 20 μm. (B) Quantification of oxidized to reduced ratio of mito-Orp1-roGFP2 in (A). Statistical significance was determined by One-way ANOVA in combination with Tukey’s test, bars represent mean ± SD. ^****^p < 0.0001, ns - not significant. (C) The redox state of the nucleus of MCF10A^ER/vSrc^ cells expressing mito-catalase and transformed with 4-OHT (7 days) was determined using fluorescence microscopy. Oxidized (λ_ex_ = 405 nM) and reduced (λ_ex_ = 488 nM) roGFP2 signals were acquired and the ratio oxidized/reduced was calculated using ImageJ (shown as a heatmap). White bars represent 20μm. (D) Quantification of oxidized to reduced ratio of NLS-Orp1-roGFP2 in (C). Statistical significance was determined by One-way ANOVA in combination with Tukey’s test, bars represent mean ± SD. ^****^p < 0.0001, ns - not significant. (E) Western blot analysis of H3.1-H3.2 and H3.3 in MCF10A^ER/vSrc^ cells expressing mito-catalase and transformed with 4-OHT (7 days). Parental cells treated with 10 nM D-Alanine for 4 h were used as positive control. (F) H3.3 / H3.1-H3.2 ratio quantification in (E). Statistical significance was determined by One-way ANOVA in combination with Tukey’s test, bars represent mean ± SD. ^****^p < 0.0001, ns - not significant. (G) EMT induction of MCF10A cells with TGFβ. Cells expressing NLS-catalase or mito-catalase were treated with 10 ng/mL TGFβ for 14 days. The levels of the Fibronectin (mesenchymal marker) and E-cadherin (epithelial marker) were determined by western blot. (H) Western blot analysis of SOX9, ZEB1 and Fibronectin in MCF10A^ER/vSrc^ cells expressing mito-catalase and transformed with 4-OHT (7 days).

### nH_2_O_2_ drives breast cancer cell chemoresistance and metastasis

Mesenchymal states are normally associated with an increase in chemoresistance as well as metastatic potential of cells in the primary tumor^23-25^. The finding that H_2_O_2_-driven chromatin remodeling is required for EMT both in a model of v-Src induced transformation as well as in response to TGFβ suggested that suppressing its activity in the nucleus could also reverse acquired resistance to chemotherapeutic drugs and inhibit the metastatic dissemination of primary xenograft tumors. We started by testing whether quenching nH_2_O_2_ reversed acquired resistance to either doxorubicin or paclitaxel. For these experiments we used two well-established murine mammary tumor cell lines derived from the aggressive MMTV.PyVT model. Although both cell lines were derived from the same model of autochthonous mammary tumorigenesis, PY230 represent more epithelial E-cadherin+, cytokeratin-8+ phenotypes while PY8119 are mesenchymal according to their N-Cadherin+, Cytokeratin-14+ and vimentin+ status. PY230^NLS-Orp1-roGFP2^ cells expressing NLS-catalase under the regulation of a tetracycline Tet-ON promoter were treated with either doxorubicin (DXR) or paclitaxel (PTX). Cell death and steady state levels of nH_2_O_2_ were simultaneously tracked for up to 14 days that included growth in drug free media for at least 2 days, drug treatment for at least 7 days before and 6 days after the induction of NLS-catalase expression. Results in Figures 5A-B indicate that PY230 cells had low steady state levels of nH_2_O_2_ prior to drug treatment. Addition of either DXR or PTX rapidly killed the bulk of cells in culture while at the same time selecting a population of persister cells displaying sustained high levels of nH_2_O_2_. Interestingly, these persister cells resumed proliferation shortly after a brief period of drug-induced dormancy (between *ca*. day 4 to day 6) unabated however by the presence of either DXR or PTX. To determine if high levels of nH_2_O_2_ were involved in the development and/or maintenance of chemoresistance, NLS-catalase expression was induced while cells were actively proliferating on day 8. As shown by fluorescence micrographs and accompanying quantification, the suppression of nH_2_O_2_ accumulation resensitized cells to the killing effects of DXR and PTX establishing a clear parallel between the accumulation of nH_2_O_2_, the development and maintenance of resistance against first line chemotherapeutic drugs. Similar results were observed when PY8119 mesenchymal cells (Figures 5C-D), although in this case basal levels of nuclear ROS were considerably higher than those observed when epithelial PY230 were used, leading to a larger number of persister cells and a significantly shorter period of drug-induced dormancy. In both cases induction of NLS-catalase downregulated ZEB1, a canonical EMT transcription factor (Figures 5E-F). Studies were also geared towards examining whether nH_2_O_2_ is necessary to enable xenograft tumor formation or unleash the metastatic potential of v-Src transformed MCF10A cells. First, we implanted wild type or NLS-catalase expressing, luciferase-labeled MCF10A^ER/vSrc^ cells in the inguinal mammary fat of immunocompromised NU/J mice. Cells were embedded in Matrigel™ containing 4-hydroxy-tamoxifen (10 μM) to induce v-Src expression. According to results shown in Figures 5G-I, only mice injected with wild-type ER-vSrc activated cells developed tumors. In the case of cells constitutively expressing NLS-catalase (and labeled with luciferase), no bioluminescence was detected even after 4 months after xenograft implantation indicating that quenching nH_2_O_2_ suppressed the ability of highly aggressive v-Src transformed cells to form tumors in mice. Given these striking observations, we decided to test the bolder idea that inducing NLS-catalase expression in established metastatic disease could halt disease progression. For these experiments we used transformed MCF10A^ER/vSrc^ cells expressing NLS-catalase under an inducible Tet-ON promoter. As expected, xenografts implanted into the inguinal mammary fat pad of female NU/J mice produced rapidly proliferating tumors that evolved to metastatic disease within three months after implantation. Surpassing our expectations, bioluminescence-based tracking of cancer cells showed activation of NLS-catalase expression led to the regression of these lesions to full remission in the majority of the mice after 1 month after doxycycline injection (Figures 5J-L). Also surprising, the effect of NLS-catalase seemed to be significantly more pronounced in metastatic lesions than on primary tumor xenografts suggesting that nH_2_O_2_ is required to sustain heterotypic metastatic colonization but may be less important to maintain tumor cell viability growing in biocompatible homotypic microenvironments. Results in Figure 5 confirmed that H_2_O_2_ has nuclear specific activities fundamentally involved in the transcriptomic and phenotypic reprogramming of tumor cells towards the acquisition of more malignant phenotypes.

**Figure 5.**
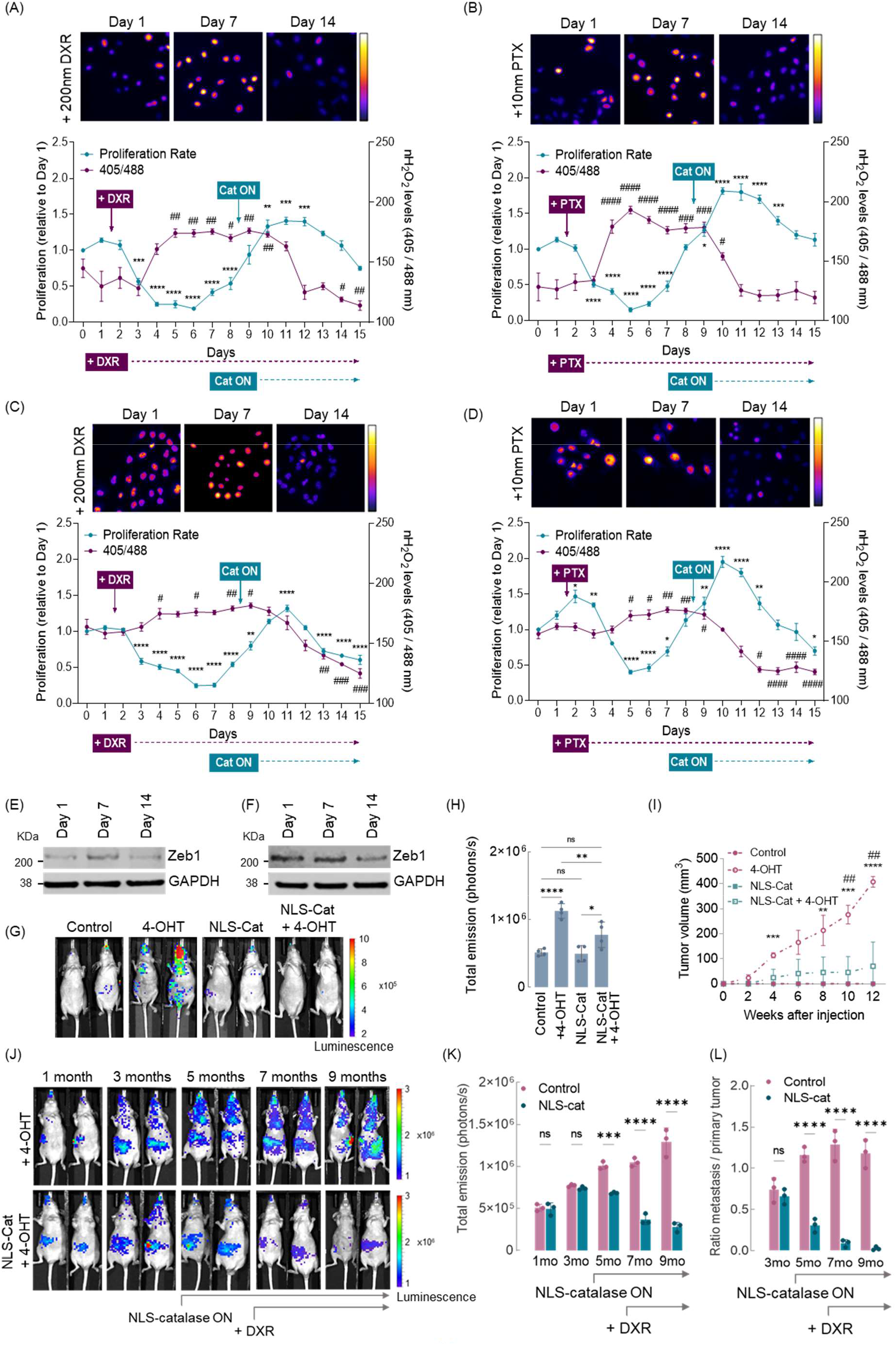
nH_2_O_2_ drives breast cancer cell chemoresistance and metastasis. (A) Cell death and the redox state of the nucleus of PY230 cells expressing Tet-ON inducible NLS-catalase and treated with 200 nM Doxorubicin (DXR). Oxidized (λ_ex_ = 405 nM) and reduced (λ_ex_ = 488 nM) roGFP2 signals were acquired and the ratio oxidized/reduced was calculated using ImageJ (shown as a heatmap). Cell death/proliferation are relative to day 1. Statistical significance of proliferation was determined by One-way ANOVA in combination with Tukey’s test, bars represent mean ± SD. ^**^p < 0.01, ^***^p < 0.001, ^****^p < 0.0001, blank - not significant. Statistical significance of the redox state of the nucleus was determined by One-way ANOVA in combination with Tukey’s test, bars represent mean ± SD. ^#^p < 0.05, ^##^p < 0.01, blank - not significant. White bars represent 50 μm. (B) Cell death and the redox state of the nucleus of PY230 cells expressing Tet-ON inducible NLS-catalase and treated with 10 nM Paclitaxel (PTX). Oxidized (λ_ex_ = 405 nM) and reduced (λ_ex_ = 488 nM) roGFP2 signals were acquired and the ratio oxidized/reduced was calculated using ImageJ (shown as a heatmap). Cell death/proliferation are relative to day 1. Statistical significance of proliferation was determined by One-way ANOVA in combination with Tukey’s test, bars represent mean ± SD. ^*^p < 0.05, ^***^p < 0.001, ^****^p < 0.0001, blank - not significant. Statistical significance of the redox state of the nucleus was determined by One-way ANOVA in combination with Tukey’s test, bars represent mean ± SD. ^#^p < 0.05, ^###^p < 0.001, ^####^p < 0.0001, blank - not significant. White bars represent 50 μm. (C) Cell death and the redox state of the nucleus of PY8119 cells expressing Tet-ON inducible NLS-catalase and treated with 200 nM Doxorubicin (DXR). Oxidized (λ_ex_ = 405 nM) and reduced (λ_ex_ = 488 nM) roGFP2 signals were acquired and the ratio oxidized/reduced was calculated using ImageJ (shown as a heatmap). Cell death/proliferation are relative to day 1. Statistical significance of proliferation was determined by One-way ANOVA in combination with Tukey’s test, bars represent mean ± SD. ^**^p < 0.01, ^****^p < 0.0001, blank - not significant. Statistical significance of the redox state of the nucleus was determined by One-way ANOVA in combination with Tukey’s test, bars represent mean ± SD. ^#^p < 0.05, ^##^p < 0.01, ^###^p < 0.001, blank - not significant. White bars represent 50 μm. (D) Cell death and the redox state of the nucleus of PY8119 cells expressing Tet-ON inducible NLS-catalase and treated with 10 nM Paclitaxel (PTX). Oxidized (λ_ex_ = 405 nM) and reduced (λ_ex_ = 488 nM) roGFP2 signals were acquired and the ratio oxidized/reduced was calculated using ImageJ (shown as a heatmap). Cell death/proliferation are relative to day 1. Statistical significance of proliferation was determined by One-way ANOVA in combination with Tukey’s test, bars represent mean ± SD. ^*^p < 0.05, ^**^p < 0.01, ^****^p < 0.0001, blank - not significant. Statistical significance of the redox state of the nucleus was determined by One-way ANOVA in combination with Tukey’s test, bars represent mean ± SD. ^#^p < 0.05, ^##^p < 0.01, ^####^p < 0.0001, blank - not significant. White bars represent 50 μm. (E) Western blot analysis of ZEB1 in PY230 cells expressing Tet-ON inducible NLS-catalase and treated with 200 nM Doxorubicin. (F) Western blot analysis of ZEB1 in PY8119 cells expressing Tet-ON inducible NLS-catalase and treated with 200 nM Doxorubicin. (G) Assessment of tumor growth and metastasis in animals injected with 4-OHT-transformed MCF10A^ER/vSrc^ cells (13 weeks after injection). The injection of MCF10A^ER/vSrc^ cells was performed subcutaneously in the mammary fat pad and tumor growth was measured weekly. Luciferase signal was detected using high exposition for 120 s. (H) Luminescent intensity quantification of (G). Statistical significance was determined by One-way ANOVA in combination with Tukey’s test, bars represent mean ± SD. ^*^p < 0.05, ^**^p < 0.01, ^****^p < 0.0001, ns - not significant. (I) Tumor volume 12 weeks after injection. Bars represent mean ± SD. Statistical significance of tumor volume was determined by One-way ANOVA in combination with Tukey’s test, bars represent mean ± SD. 4-OHT x Control: ^**^p < 0.01, ^**^p < 0.001, ^****^p < 0.0001, blank - not significant. 4-OHT x NLS-Cat + 4-OHT: ^##^p < 0.01, blank - not significant. (J) Assessment of tumor growth and metastases in animals injected with MCF10A^ER/vSrc^ cells expressing Tet-ON inducible NLS-catalase and treated with 5 mg/kg Doxorubicin (DXR). The injection of MCF10A^ER/vSrc^ cells was performed subcutaneously in the mammary fat pad and tumor growth and metastasis were measured monthly. The induction of NLS-catalase occurred after the fourth month and the treatment with DXR was performed in a weekly basis after the fifth month. Luciferase signal was detected using high exposition for 120 s. (K) Luminescent intensity quantification of (J). Statistical significance was determined by One-way ANOVA in combination with Tukey’s test, bars represent mean ± SD. ^***^p < 0.001, ^****^p < 0.0001, ns - not significant. (L) Ratio metastasis / primary tumor of (J). Statistical significance was determined by One-way ANOVA in combination with Tukey’s test, bars represent mean ± SD. ^****^p < 0.0001, ns - not significant.

## DISCUSSION

In eukaryotes, the progressive accumulation of H3.3 over H3.1 represents an indelible hallmark of chronologic aging^26^. Another hallmark of aging is progressive metabolic/mitochondrial dysfunction invariably associated with an increase in the production and/or a decrease in the detoxification of ROS^27^. Both an increase in ROS and the exchange of histone variants have also been shown to occur in cancer and to be involved in its progression towards metastasis^9^. Therefore, we investigated the possibility that oxidation-induced H3.1 loss might represent a new mechanistic link between metabolic decline and epigenetic reprogramming associated with the evolution of cancer, a disease that disproportionally affects the elderly. The idea that mitochondria ROS might influence epigenetic reprogramming was also reinforced by findings that mitochondria and the nucleus are connected. In 2020, Desai et al.^28^, showed that mitochondria regulated gene transcription, at least in part, via the transfer of ROS via tethers that connect the organelles. Consistently, we found that most transformed cancer cells displaying more aggressive behavior and higher degrees of mitochondrial dysfunction existed in states characterized by higher steady state levels of nuclear ROS. To recapitulate this effect and understand it is consequences for the regulation of gene expression we used the direct delivery of H_2_O_2_ by an engineered D-amino acid oxidase system fused to a nuclear localization sequence (NLS). Delivery of ROS to the nucleus caused loss of H3.1 changing the H3.1/H3.3 ration ultimately promoting chromatin decompaction and the expression of genes normally silenced by localization into heterochromatic domains. These include EMT genes. As shown before H3.1 replacement by H3.3 preceded and was required to license EMT either induced by TGFβ or triggered directly via the oxidation of H3.1-Cys96. Mutation of the redox active Cys96 residue conferred resistance to ROS-induced as well as TGFβ-induced H3.1 loss and EMT confirming that Cys96 oxidation as a critical event in H3.1 destabilization and chromatin remodeling associated with cellular plasticity acquisition. Consistently, expressing mutant H3.1 lacking Cys96 or quenching of ROS with catalase directed to the nucleus (NLS-catalase) suppressed H3 variant exchange induced by ROS as well as ROS-induced EMT gene expression. *In vivo*, suppressing nuclear ROS inhibited tumorigenesis, tumor growth and strongly suppressed metastasis. Surprisingly, suppression of nuclear ROS in established metastatic lesions drove them into remission indicating that nuclear ROS are not only critical to enable metastatic colonization but also to support the viability and growth of the metastatic lesion at heterotypic organs. Hence, by demonstrating that an increase in nuclear ROS structurally regulates EMT gene accessibility via oxidation of redox sensitive H3.1 our results mechanistically connect several hallmark features common to aging, metabolic changes occurring in aging and cancer including: 1) the enrichment of H3.3 over H3.1; 2) an increase in steady state ROS; 3) epigenetic activation of EMT and stemness genes shown here to be induced by and dependent on mitochondrial H_2_O_2_ reaching the nucleus; and 4) increased cellular resistance to chemotherapeutics and metastatic potential that could be, to a large extent, backtracked to epigenetic reprogramming executed directly by H_2_O_2_. Taken together our results suggest that mitochondrial-ROS driven chromatin remodeling is part of a cell survival program activated by acute metabolic stress to expand the range of transcriptional possibilities cells can explore. However, in cancer cells, the expansion of the cells’ transcriptional potential via oxidative chromatin remodeling supports clonal plasticity particularly under the selective pressure of chemotherapy, a problem difficult to circumvent clinically. Therefore, we believe that there is significant therapeutic potential in the manipulation of mechanisms based on redox epigenetics for the development of a novel class of treatments for cancer and, potentially other diseases of aging.

## MATERIALS AND METHODS

### Cell culture

Cell lines were grown in 5% CO_2_ at 37 °C. MCF10A^ER/vSrc^ cell line was a generous gift from Kevin Struhl, Harvard University, Cambridge, MA. HEK293, MCF7, BT474, BT20 and MDA-MB231 cells were obtained from American Type Culture Collection (ATCC). MCF10A^ER/vSrc^ cells were propagated in DMEM/F12 (1:1) (Gibco, #21041025) supplemented with 5% charcoal stripped fetal bovine serum (Sigma Aldrich, #F6765), 20 ng/ml EGF (Sigma Aldrich, #SRP3027), 10 μg/ml insulin (Sigma Aldrich, #I3536), 0.5 μg/ml hydrocortisone (Sigma Aldrich, #H0135), 0.1 μg/ml cholera toxin (Sigma Aldrich, #C8052), and 1% penicillin-streptomycin. MCF7, BT474, BT20 and MDA-MB231 cells were cultured in DMEM supplemented with 10% fetal bovine serum (Sigma Aldrich, #F2442) and 1% penicillin-streptomycin. Cells were regularly tested for mycoplasma. For malignant transformation, MCF10AER/vSrc cells were treated with 1μM 4-hydroxytamoxifen (4-OHT) (Sigma, #H7904) for 7 days.

### Plasmids, lentivirus and viral transduction

The eukaryotic expression vectors encoding nuclear localization tagged proteins were generated by inserting polymerase chain reaction (PCR)–amplified fragments into the adequate backbone vector. NLS-Orp1-roGFP2 and mito-Orp1-roGFP2 (puromycin as selection mark) were obtained from cloning the c-myc sequence or a MTS sequence respectively into the Orp1-roGFP2 vector (Addgene, #64993). NLS-DAO plasmid (geneticin as selection mark) was a generous gift from Thomas Michel, Harvard University. NLS-catalase and TET-ON NLS-catalase plasmids were obtained from cloning the catalase and c-myc sequences into a hygromycin-containing backbone vector (Addgene, #17446). H3.1 C96S-Flag (blasticidin as selection mark) was obtained from cloning the H3.1 C96S and Flag sequences into a backbone vector (Addgene, #17445). Fast-FUCCI plasmid was obtained from Addgene (#86849). Luciferase plasmid (blasticidin as selection mark) was obtained from cloning the luciferase sequence into a blasticidin-containing backbone vector (Addgene, #17445). HEK293 cells were transfected at 70 to 80% confluency using Lipofectamine 3000 (Thermo Fisher Scientific, #L3000015) and incubated at 37°C for 2 days. Supernatant was collected, filtered and used to transduce MCF10A^ER/vSrc^, MCF7 and MDA-MB231 cells. After transduction (2 days), cells were selected with appropriate antibiotics.

### Oxidized and reduced Orp1-roGFP2 detection

Cells (untreated, treated with 10 nM D-Alanine for 4 h or 500 μM H_2_O_2_ for 30 min) were imaged in Nikon W1 Dual Cam Spinning Disk Confocal automated microscope or in Lionheart FX (Biotek). Orp1-roGFP2 was excited sequentially at 405 and 488 nm and the emission was recorded at 525 nm. The generated images were analyzed using FIJI ImageJ (1.53c). Each channel was adjusted using the default FIJI ImageJ threshold algorithm. Only the nuclear area was considered for the analysis. Oxidation levels were then determined on a pixel by pixel basis dividing the signal emitted for each channel (405 nm / 488 nm). Final ratiometric values were obtained using the “Analyzing particles” tool and a heatmap was created for visualization.

### 8-oxo-dG detection

MCF10A^ER/vSrc^ cells were plated in black bottom 96-well plates and treated with 10 nM D-Alanine for 4 h or 500 μM H_2_O_2_ for 30 min. Subsequently, cells were fixed with 4% formaldehyde solution (Sigma Aldrich, #F8775) for 15 min at room temperature and permeabilized with ethanol (Sigma Aldrich, #E7023) for 1 min. Then, cells were washed with PBS pH 7.4 and blocked with 5% (w/v) bovine serum albumin (BSA) (Sigma Aldrich, #A9648) for 2 h at 4 °C and finally stained with anti-8-oxo-dG (Trevigen, #4354-MC-050), followed by incubation with secondary antibody Alexa647 (Invitrogen #A28181). Images were collected under automatic microscope Lionheart FX (Biotek).

### Comet assay

Comet assay was performed using the Comet Assay Kit according to the instructions provided by the manufacturer (Trevigen, #4250-050-K). MCF10A^ER/vSrc^ cells were seeded in 6-well plates and treated with 10 nM D-Alanine for 4 h or 500 μM H_2_O_2_ for 30 min. Cells were then washed, detached and resuspended in PBS solution. The cell suspension was mixed with LM-agarose at a ratio of 1:10 (v/v), transferred onto the slides, and lysed at 4 °C. The slides were treated with an alkaline unwinding solution for 60 min and placed on a horizontal electrophoresis unit filled with fresh buffer alkaline unwinding solution. Electrophoresis was performed at 20 V for 35 min at 4 °C in the dark and subsequentially staining. Images were collected under automatic microscope Lionheart FX (Biotek). The tail moment and % DNA in tail were analyzed using the Open Comet plug-in in FIJI ImageJ (1.53c).

### Amplex red assay

H_2_O_2_ was measured using an Amplex Red Hydrogen Peroxide/Peroxidase Assay Kit (Thermo Fisher Scientific, A22188), according to the manufacturer protocol. MCF10A^ER/vSrc^ cells were plated in black bottom 96-well plates and treated with different concentrations of D-Alanine (10 nM, 10 μM, 100 μM, and 1 mM). 50 μL Amplex Red reagent with 0.2 units/mL horseradish peroxidase was added to samples and fluorescence was measured for 2 h at 37 °C (λ_ex_ = 530 nm, λ_em_ = 590 nm). The mean reading was plotted on a simultaneously prepared H_2_O_2_ standard curve.

### Western blot analysis

Cells were lysed in cold RIPA buffer (Thermo Fisher Scientific, #89901) supplemented with protease inhibitor cocktail (Thermo Fisher Scientific, #87785). Equal amounts of total protein (30 to 50 μg) were separated in a 4–12% Bis-Tris polyacrylamide gels (Invitrogen, #NP03354) and transferred to a 0.22 μm nitrocellulose membrane (BIO-RAD, #1620112). Membranes were incubated with appropriate primary antibodies (phospho-ATM, 1:1000, Cell Signaling Technology, #13050; E-Cadherin, 1:1000, Cell Signaling Technology, #3195; Fibronectin, 1:1000, Cell Signaling Technology, #26836; Flag, 1:1000, Cell Signaling, #14793; GAPDH, 1:1000, Cell Signaling Technology, #97166; H2AX, 1:1000, Cell Signaling Technology, #7631; phospho-H2AX, 1:1000, Cell Signaling Technology, #8032; H3.1-H3.2, 1:1000, Millipore, #09-838; H3.3, 1:1000, Millipore, #ABE154; PKM1/2, 1:1000, Cell Signaling Technology, #3106; SOX9, 1:1000, Cell Signaling Technology, #82630; TBP, 1:1000, Cell Signaling Technology, #44059; ZEB1, 1:1000, Cell Signaling Technology, #3396) and species-specific antibodies (IRDye Goat anti-Mouse, 1:10000, LI-COR, #925-68070; IRDye Goat anti-Rabbit, 1:10000, LI-COR, #9263221). Signal was detected using an Odyssey FC (LI-COR) imaging station.

### RNA extraction and RT-qPCR

Total RNA was extracted from MCF10A^ER/vSrc^ cells (Control and treated with 10 nM D-Alanine for 4 h or 24 h) using RNeasy Kit (Qiagen, #74104) according to the manufacturer’s instructions. cDNA was synthesized using High Capacity cDNA Reverse Transcription kit (Applied Biosystem, #4368814). Quantitative PCR was performed on an Applied Biosystems QuantStudio 6 Flex Real-Time PCR System using Fast SYBR Green Master Mix (Applied Biosystems, #4385612). The relative gene expression was calculated using the 2^−ΔΔCt^ method with GAPDH as endogenous control for normalization.

### RNA-Seq

RNA was isolated from MCF10A^ER/vSrc^ cells as described above. Each sample was treated with 10 nM D-Alanine for 4 h or 24 h. Sequencing libraries were generated using Illumina Novaseq platform. Sample quality, library complexity, and alignment statistics were checked using an established pipeline in our institution.

### RNA-Seq data analysis

All RNA-Seq analysis were performed with 5 biological replicates. The sequencing reads were aligned against the reference human genome hg19. Differentially expressed genes (DEG) were identified using EdgeR in control and 10nM D-Alanine (4 h or 24 h) with cutoff ≥ 1.5-fold increase in expression compared to control, by pairwise approach. Heatmap was generated using normalized expression levels of differentially expressed genes in pairwise comparison using GraphPad Prism 9. GSEA (Hallmark genes analysis) was performed using the 10 nM for 4 h x Control data set.

### Chromatin immunoprecipitation

Approximately 5 × 10^6^ MCF10A^ER/vSrc^ cells (Control and treated with 10 nM D-Alanine for 4 h) were fixed with 1% methanol-free formaldehyde solution (Thermo Scientific, #28908) for 10 min at RT. Then, reaction was quenched with addition of 125 mM glycine (Sigma Aldrich, #G8898) for 5 min. Chromatin isolation and immunoprecipitation were carried out using Magna ChIP A/G kit (Millipore, #1710085). Chromatin was isolated from nuclei and sonicated in Diagenode Bioruptor Pico for 10 cycles of 30 sec with intervals of 30 sec at 4 °C to generate DNA fragments with size range of 200 to 600 bp. Sheared chromatin equivalent to 2 × 10^6^ cells was diluted 1:10 in ChIP Dilution Buffer and protease inhibitor cocktail and incubated for 16 h with 20 μL Magnetic Protein A/G Beads (Millipore #CS204457) and 5 μL antibodies. The antibodies used were H3.1-H3.2 (Millipore, #09-838), H3.3 (Millipore, #ABE154), Flag (Cell Signaling, #14793), HA (Cell Signaling, #3724) or Normal Rabbit IgG (Cell Signaling, #2729). Immunoprecipitated DNA fragments were eluted and purified according to manufacturer’s instructions.

### ChIP-Seq

Immunoprecipitated DNA from MCF10A^ER/vSrc^ (Control and treated with 10 nM D-Alanine for 4 h) was tested for concentration, integrity and purity. Concentration was detected by fluorometer or Microplate Reader (Qubit Fluorometer, Invitrogen). Sample integrity and purity were detected by Agilent Technologies 2100 bioanalyzer. ChIP’s DNA was subjected to end-repair and then was 3’ adenylated. Adaptors were ligated to the ends of these 3’ adenylated fragments. DNA fragments were amplified with adaptors from previous step. PCR products were purified and selected with the Agencourt AMPure XP-Medium kit. The double stranded PCR products were heat denatured and circularized by the splint oligo sequence and the single strand circle DNA (ssCir DNA) were formatted as the final library. Library was qualified by Qubit ssDNA kit amplified to make DNA nanoball (DNB), which had more than 300 copies of one molecular. The DNBs were loaded into the patterned nanoarray and single end 50 bases reads were generated in the way of sequenced by combinatorial Probe-Anchor Synthesis (cPAS).

### ChIP-Seq data analysis

The quality of reads was evaluated using FastQC. Reads were trimmed from the 3’ ends using cutadpat and aligned using Bowtie 2 (version 2.2.9) with default parameters to the human genome (hg38). Reads were then analyzed with Homer (version 4.11) to call H3.1-H3.2 and H3.3 binding regions, identify regions of differential binding and generate IGV read density tracks.

### ChIP-qPCR

ChIP products were analyzed by qPCR using Fast SYBR Green (Applied Biosystems #4385612) on a Quant Studio 6 Flex PCR system (Applied Biosystems). Each ChIP sample was normalized by its respective 1% input’s adjusted Ct (ΔCt), where

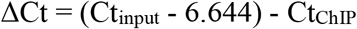

To express ChIP enrichment in percentage of input, the formula used was

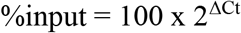

Each experiment was performed in three independent biologic replicates. Error bars represent standard deviation of replicates. Statistical analysis was performed by two-way ANOVA corrected for multiple comparisons using Turkey method and confidence interval level of 95%.

### Transmission Electron Microscopy

MCF10A^ER/vSrc^ cells (Control and treated with 10 nM D-Alanine for 4 h) were fixed in mixture of 2.5% glutaraldehyde and 2% paraformaldehyde in 0.1 M cacodylate buffer overnight at 4 °C. After post-fixation in 1% osmium tetroxide and 3% uranyl acetate, cells were dehydrated in series of ethanol, embedded in Epon resin and polymerized for 48 h in 60 °C. Then ultrathin sections were made using Ultracut UC7 Ultramicrotome (Leica Microsystems) and contrasted with 3% uranyl acetate and Reynolds’s lead citrate. Samples were imaged using a FEI Tecnai Spirit G2 transmission electron microscope (FEI Company, Hillsboro, OR) operated at 80 kV. Images were captured by Eagle 4k HR 200kV CCD camera. Images were processed using FIJI ImageJ (1.53c). The percentage of heterochromatin in at least 20 nuclei was determined using Trainable Weka Segmentation plug-in.

### EMT induction with TGFβ

MCF10A cells with or without expression of NLS-catalase or mito-Catalase were treated with 10 ng/mL TGFβ (Sigma, #H8541) every 2 days for a total of 14 days. Cells were then prepared for Western blot analysis and total protein was incubated with E-Cadherin (1:1000, Cell Signaling Technology, #3195), Fibronectin (1:1000, Cell Signaling Technology, #26836) and GAPDH (1:1000, Cell Signaling Technology, #97166) followed by species-specific antibodies incubation (IRDye Goat anti-Mouse, 1:10000, LI-COR, #925-68070; IRDye Goat anti-Rabbit, 1:10000, LI-COR, #9263221). Signal was detected using an Odyssey FC (LI-COR) imaging station.

### Chromatin accessibility assay

Chromatin accessibility was analyzed using Chromatin Accessibility Assay Kit (Abcam, #ab185901). Cells treated with D-Alanine for different timepoints were lysed and chromatin was extracted. Chromatin was thereafter digested using a nuclease mix and the purified DNA was analyzed by qPCR using specific primers for Fibronectin, SOX9, and ZEB1. Results were calculated by fold enrichment using a ratio of amplification efficiency of nuclease-treated samples over that of untreated nuclease-free control samples as suggested in the protocol.

### Drug Resistance Studies

PY230 and PY8119 cells were transduced with TET-ON NLS-catalase and treated with 200 nM Doxorubicin (DXR) or 10 nM Paclitaxel (PTX). The expression of NLS-catalase was performed by treating the cells with 100 ng/mL doxycycline every two days. The redox state of the nucleus and cell death/proliferation were measured daily. Cells were collected in specific time points for western blot analysis of EMT markers (ZEB1).

### Animal studies

All mouse experimentation was conducted in accordance with standard operating procedures approved by the Institutional Animal Care and Use Committee of our Institution. NU/J mice were acquired from The Jackson Laboratory (#002019). For subcutaneous xenograft, 25 μL medium containing 1 × 10^6^ luciferase-expressing MCF10A^ER/vSrc^ cells (control or 4-OHT transformed, with or without NLS-catalase expression) were mixed with 75 μL Matrigel matrix (BD Bioscience, #356237) containing 10 μM 4-OHT and injected into the inguinal mammary fat pad of the female mice at 6 to 8 weeks of age. The diameters of tumor were measured every two weeks with a digital caliper, and the tumor volume in cubic millimeters was calculated using the formula (length × width^2^/2). Imaging of grafted tumors was performed using an IVIS in vivo imager (LAGO) after intraperitoneal injection of RediJect D-Luciferin Bioluminescent Substrate (PerkinElmer, #770504) 13 weeks after injection. Luminescent intensity was acquired using high exposition for 120s and total emission (photons/s) was calculated. Similarly, NU/J animals were injected with transformed MCF10A^ER/vSrc^ cells expressing NLS-catalase under an inducible Tet-ON promoter. The expression of NLS-catalase was induced using doxycycline. Animals were treated with 5 mg/kg Doxorubicin (DXR) every two weeks after the fifth month of the injection. Imaging of grafted tumors was performed monthly using an IVIS in vivo imager (LAGO) after intraperitoneal injection of RediJect D-Luciferin Bioluminescent Substrate (PerkinElmer, #770504).

### Statistical analysis

Statistical analysis was performed with GraphPad Prism v9 (GraphPad Software Inc.). Data with a normal distribution are expressed as the mean ± SD. A value of *p* < 0.05 was considered significant.

## AUTHORS’ CONTRIBUTION

FRP, BNG, SR, VB and MGB conceptualized the study. FRP, FTO, DRC, KP, AM, YH and JMD performed experiments. FRP, FTO, DRC and KP analyzed data. FRP, CMF, DS, BNG, SR, VB and MGB interpreted data. FRP, FTO and KP prepared figures. FRP, BNG and MGB wrote the paper.

## DATA AND MATERIALS AVAILABILITY

RNA-Seq and ChIP-Seq data will be available in NCBI GEO. All other data and materials are available in the paper and supplementary materials.

