## Supplementary Data for "H3.1^Cys96^ oxidation by mitochondrial ROS promotes chromatin remodeling, breast cancer progression to metastasis and multi-drug resistance"

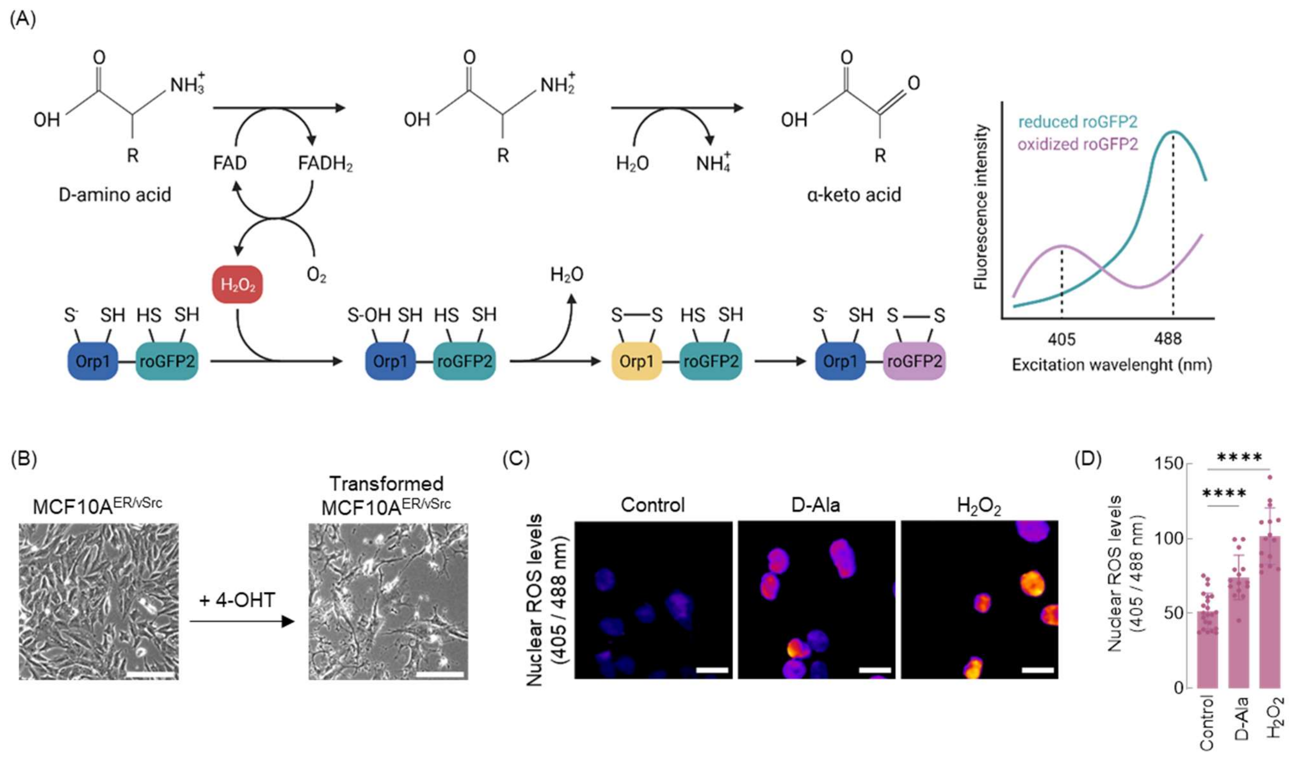


**Supplementary Figure 1. NLS-DAO / NLS-Orp1-roGFP model.**

(A) Model of coupled nuclear $H_2O_2$ production and sensing. We generated a model for nuclear production of $H_2O_2$ using D-amino acid oxidase (DAO), an enzyme that produces $H_2O_2$ as a byproduct of D-amino acids oxidation. To detect nuclear DAO-derived $H_2O_2$ we expressed the nuclear redox sensor Orp1-roGFP2. Orp1 is oxidized by $H_2O_2$ and consequently oxidizes roGFP2. Reduced (405 nm excitation) and oxidized (488 nm excitation) roGFP2 are detected using fluorescence/confocal microscopy.

(B) Epithelial MCF10A$^{ER/vSrc}$ cells are transformed to mesenchymal and tumorigenic phenotypes under induction of vSrc oncogene with 4-OHT. White bars represent 200 mm.

(C) MCF10A$^{ER/vSrc}$ cells were treated with 10 nM D-Alanine for 4 h and exogenous $H_2O_2$ (500 µm for 30 min). The redox state of the nucleus was determined using confocal microscopy. Oxidized and reduced Orp1-roGFP2 signals were acquired (405 nm and 488 nm excitation respectively, 525 nm emission) and the ratio oxidized / reduced is shown as a heatmap.

(D) Quantification of oxidized (405 nm excitation) to reduced (488 nm excitation) ratio of Orp1-roGFP2 in (C). Statistical significance was determined by One-way ANOVA in combination with Tukey's test, bars represent mean ± SD. $^{****}p < 0.0001$.

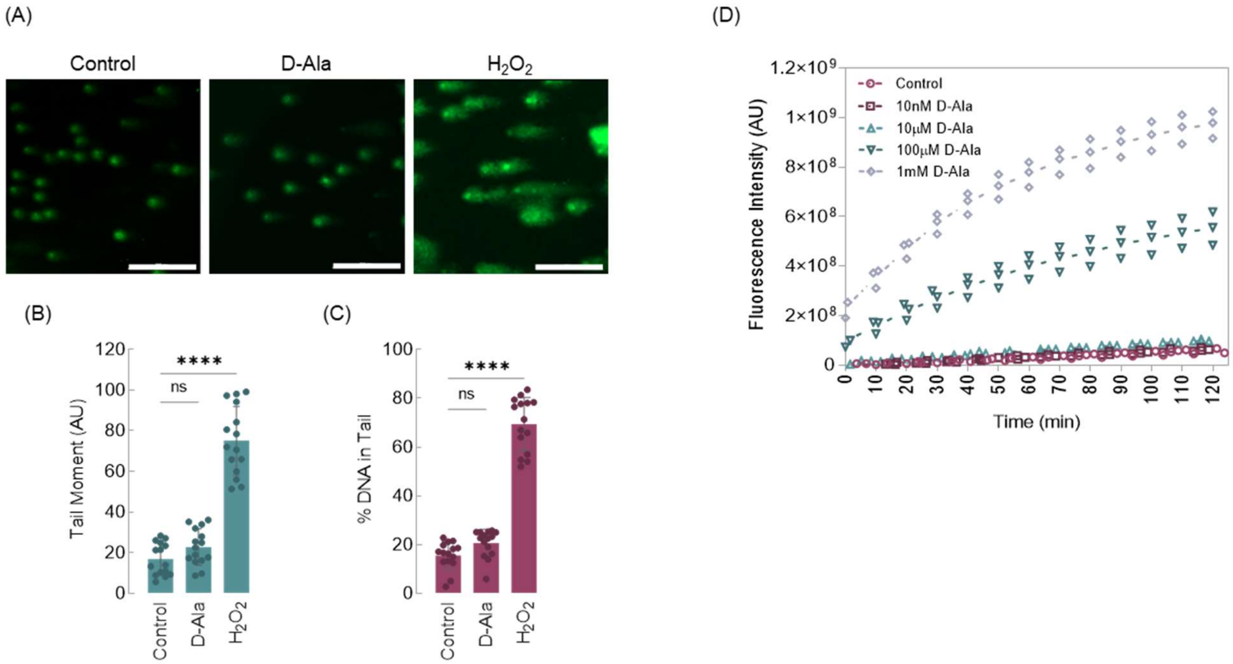


**Supplementary Figure 2. Assessment of DNA damage and extracellular $H_2O_2$ levels in the study model.**

(A) Comet assay analysis of DNA strand breaks in MCF10A$^{ER/vSrc}$ cells treated with 10 nM D-Alanine for 4 h and exogenous $H_2O_2$ (500 µm for 30 min). White bars represent 100 mm.

(B) Quantification of DNA damage by tail moment of comet assay. Statistical significance was determined by One-way ANOVA in combination with Tukey's test, bars represent mean ± SD. $^{****}p < 0.0001$, ns - not significant.

(C) % DNA in comet tails. Statistical significance was determined by One-way ANOVA in combination with Tukey's test, bars represent mean ± SD. $^{****}p < 0.0001$, ns - not significant.

(D) Extracellular $H_2O_2$ detection. MCF10A$^{ER/vSrc}$ cells were treated with different concentrations of D-Alanine (10 nM, 10 mM, 100 mM and 1 mM) and the levels of $H_2O_2$ released to extracellular milieu was measured using Amplex red.
